# Generalization and search in risky environments

**DOI:** 10.1101/227322

**Authors:** Eric Schulz, Charley M. Wu, Quentin J. M. Huys, Andreas Krause, Maarten Speekenbrink

## Abstract

How do people pursue rewards in risky environments, where some outcomes should be avoided at all costs? We investigate how participant search for spatially correlated rewards in scenarios where one must avoid sampling rewards below a given threshold. This requires not only the balancing of exploration and exploitation, but also reasoning about how to avoid potentially risky areas of the search space. Within risky versions of the spatially correlated multi-armed bandit task, we show that participants’ behavior is aligned well with a Gaussian process function learning algorithm, which chooses points based on a safe optimization routine. Moreover, using leave-one-block-out cross-validation, we find that participants adapt their sampling behavior to the riskiness of the task, although the underlying function learning mechanism remains relatively unchanged. These results show that participants can adapt their search behavior to the adversity of the environment and enrich our understanding of adaptive behavior in the face of risk and uncertainty.

## Introduction

Your phone rings. It is your parents. They are on their way for a surprise visit. You can hear the engine of their car running as you talk to them. They will arrive in a few hours and ask if you could prepare something for dinner. Your mum jokes that they already had beans on toast for lunch. Aiming to amaze them with a unique culinary experience, you decide to prepare something extraordinary, something they have never eaten before. As you open your fridge and kitchen cupboards, you find a plethora of ingredients at your disposal. In your head, you go through different combinations of ingredients, imagining how their taste combines and interacts to produce a—hopefully memorable—culinary experience. You have enough time to try out some combinations, experiencing the resulting taste, and thereby learning about the effects of unusual combinations and methods of preparation. While you can be adventurous, you need to be sure to avoid certain options; you cannot risk trying inedible, poisonous, or otherwise disappointing dishes.

This scenario is an example of a multi-armed bandit task (Robbins, 1985; Srinivas, Krause, Kakade, & Seeger, 2009; Steyvers, Lee, & Wagenmakers, 2009), where there are a number of options or “arms” of the bandit (e.g., the possible dishes) which lead to initially unknown and stochastic outcomes or rewards (e.g., the taste of the dish), that are related to a set of features (e.g., the ingredients, the method of preparation, and so forth). Through experience, you can learn a function which maps features to rewards, and use this knowledge to maximize the overall rewards gained over repeated plays of the bandit. A key challenge for optimal behavior in such tasks is framed by the exploration-exploitation dilemma (Gittins, 1979; Laureiro-Martínez, Brusoni, & Zollo, 2010): should you choose an option that you know will likely lead to a high reward, or try an unknown option to experience its outcome and thereby learn more about the function mapping features to rewards, increasing the chances of gaining even higher rewards in the future?

Single-mindedly focusing on optimizing outcomes is frequently ill-advised as there might be further constraints which one has to take into account. For example, you may need to avoid a particularly bad outcome (e.g., poisonous food) at all cost. In order to satisfy this constraint, you should only explore options that—while uncertain—are likely to be “safe”. Such restricted exploration-exploitation problems are indeed common in daily life, from choosing which restaurant to visit (avoid food poisoning), where to buy a second-hand car (avoid buying a lemon), to finding the shortest route home (avoid dangerous terrain). In our previous research on human behavior in contextual (Schulz, Konstantinidis, & Speekenbrink, 2017) and spatially-correlated multi-armed bandits (Wu, Schulz, Speekenbrink, Nelson, & Meder, 2018), we found that human behavior in the search for rewards without constraints can be robustly described by a combination of a universal function learning mechanism and a decision strategy which explicitly balances an option’s expected reward and its attached uncertainty. The function learning mechanism was formalized as Gaussian process regression, which is a form of non-parametric Bayesian regression that adapts its complexity to the data at hand (Griffiths, Lucas, Williams, & Kalish, 2009; Rasmussen, 2006), while the decision strategy was formalized as upper confidence bound sampling strategy (UCB; Auer, 2002).

In the present study, we expand on our previous work by introducing scenarios with additional constraints: unsafe options—defined as options which produce outputs below a given threshold—which should be avoided at all costs. We assess how people behave when they have to maximize accumulated rewards while also avoiding (momentary) outcomes below the threshold. The task is presented as a spatially correlated multi-armed bandit in which participants choose an input, and then observe and accrue the output of an underlying function which maps spatial locations to expected rewards. In two experiments with a uni- and bivariate spatially-correlated multi-armed bandit, we find that participants adapt their exploration-exploitation strategy to the additional constraints of risky situations, but utilize the same underlying learning mechanism.

From a computational perspective^1^, the task of maximizing rewards while avoiding unsafe inputs can best be solved by a safe optimization algorithm (Sui, Gotovos, Burdick, & Krause, 2015). This algorithm tries to find points that are likely to be safe, and then expands on the set of safe points while also attempting to optimize the underlying function. When analyzing how humans choose from the three sets of points identified by a safe optimization algorithm (i.e., safe, expanding, and maximizing points), we find that choices are strongly influenced by a tendency to stay safe.

From an algorithmic perspective, it is possible to predict individual participant choices by combining different models of learning with multiple decision strategies, and then perform model comparison using out-of-sample prediction accuracy. Whereas the estimated parameters of the learning model remain relatively unchanged, we find that participants seem to adapt their decision strategy to whether or not they need to avoid unsafe outcomes, as predicted by a decision strategy that focuses on staying safe.

Our results point to the relevance of safe reinforcement learning (Berkenkamp, Turchetta, Schoellig, & Krause, 2017) for explaining human behavior in naturalistic tasks and enrich our notion of how people strategically adapt their behavior to the risk and uncertainty of the environment. In particular, whereas the way in which people generalize over different options remains similar across different riskiness conditions, they tend to adapt their sampling strategy by focusing more on safe actions as the situations become more risky.

## General task description

We use a variant of the spatially correlated multi-armed bandit (Wu, Schulz, Speekenbrink, Nelson, & Meder, 2017), where the rewards of each option (i.e., arm) are correlated according to how close they are to each other. Intuitively, nearby arms tend to have similar rewards, with the level of similarity decreasing over larger distances. The options are either univariate input values placed along a line, or bivariate input values placed on a grid. Each discretized input value represents a playable arm of the bandit. Traditionally, the goal in such tasks is to maximize cumulative payoffs by sequentially choosing one of the *N*-arms of the bandit that stochastically generate rewards (Lai & Robbins, 1985; Steyvers et al., 2009), with learning happening independently for each arm (e.g., through associative learning). In our case, because proximate arms generate similar rewards due to the spatial correlations, there is the opportunity to form inductive beliefs about the rewards of untried options by learning the function that maps the spatial location of options to their rewards. This task allows us to study how people generalize their experience to generate beliefs about novel options, and how this process influences their search behavior (Wu et al., 2018).

Importantly, we add a constraint such that participants need to avoid rewards below a given threshold. If participants obtain rewards above the threshold, they collect the reward and continue to the next trial where they are again asked to choose an input. If they obtain a reward below the threshold, they forgo the reward, end their round, and lose the opportunity to collect further rewards within the current round.

The additional requirement of avoiding unsafe options in the Risky Spatially Correlated Bandit makes generalization even more important, as it now serves to identify not only highly rewarding options, but also unsafe options. In contrast to unconstrained spatially correlated multi-armed bandits, where good performance does not require accurate knowledge of the function in regions of low rewards, our risky version requires people to learn about both regions of low and high reward.

## Function learning as model of generalization

We assume that generalization within spatially correlated multi-armed bandits can be described as a function learning mechanism that learns a function mapping the spatial context of each arm to expectations of reward. We use Gaussian process regression (Rasmussen, 2006; Schulz, Speekenbrink, & Krause, 2016) as an expressive model of human function learning. Gaussian process regression is a non-parametric Bayesian approach towards function learning which can perform generalization by making inductive inferences about unobserved outcomes. In past research we found that Gaussian process regression captures the inductive biases of human participants in a variety of explicit function learning tasks (Schulz, Tenenbaum, Duvenaud, Speekenbrink, & Gershman, 2016), and provides an accurate description of human generalization in contextual and spatially correlated multi-armed bandits without the presence of unsafe outcomes (Schulz et al., 2017; Wu et al., 2018).

Gaussian process regression integrates both rule-based and similarity-based approaches towards function learning and has originally been proposed as a rational model of human function learning by Lucas, Griffiths, Williams, and Kalish (2015). Here, we use Gaussian process regression both as a rational model, and as a component in our models which describe behavior on an algorithmic level. Thus, we use an approach that bridges the gap between two levels of descriptions (Griffiths, Lieder, & Goodman, 2015; Griffiths, Vul, & Sanborn, 2012).

To categorize participant choices from a computational level, we assess the correspondence between input points preferred by a Gaussian process safe optimization algorithm and those preferred by the participants in our experiments. To determine the model which describes behavior on the algorithmic level, we combine Gaussian process regression with different decision strategies, some of which are risk-averse and some of which are not. We then use cross validation to compare the resulting models with other models that do not apply generalization.

### Gaussian process function learning

A Gaussian process defines a distribution over functions (see Rasmussen, 2006; Schulz, Speekenbrink, & Krause, 2016, for an introduction). Let 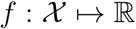 denote a function over input space 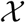 (i.e., options or arms) that maps to real-valued scalar outputs (i.e., rewards). The function is assumed to be a random draw from a Gaussian process:

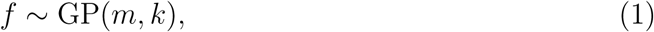

where *m* is a mean function specifying the expected output of the function given input x, and *k* is a kernel (or covariance) function specifying the covariance between outputs:

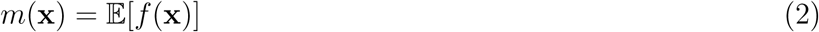

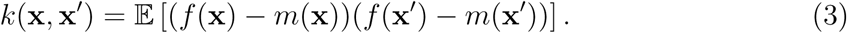

Intuitively, the kernel encodes an inductive bias about the function’s expected smoothness. We follow standard conventions and set *m*(*x*) = 0.

Conditional on observed data 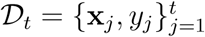, where 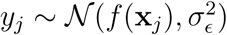 noise-corrupted draw from the underlying function (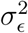 is the noise variance), the posterior predictive distribution of the function value for a new input **x**_*_ is Gaussian with mean and variance given by:

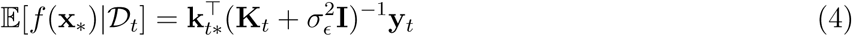

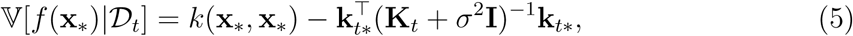

where **y***_t_* = [*y*_1_,…, *y_t_*]^⊤^, **K***_t_* is the *t* × *t* matrix of covariances evaluated at each pair of observed inputs, and *k_t_***_*_** = [*k*(**x**_1_, **x_*_**),…, *k*(x*_t_*, x**_*_**)] is the covariance between each observed input and the new input **x**_*_.

A common choice of kernel function is the radial basis function kernel

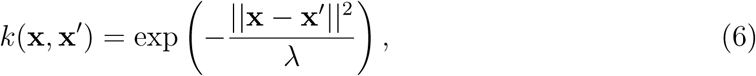

where the length-scale λ governs how quickly correlations between points **x** and **x**′ decay towards zero as their distance increases^2^.

## Rational model

Given a learned representation of a function at time *t*, this knowledge can be used to choose the next input at time *t* + 1 in a close-to-rational way. This is done through a *decision strategy* that takes the predicted mean *μ*(**x**) and uncertainty *σ*(**x**) for each input, and produces a criterion governing which input to choose next in order to balance exploration and exploitation (Brochu, Cora, & De Freitas, 2010; Schulz, Speekenbrink, & Krause, 2016).

A strategy that can cope with the additional requirement to avoid outcomes below a threshold in a close-to-rational way, is the *safe optimization algorithm* put forward by Sui et al. (2015). This algorithm uses Gaussian process regression to form beliefs about the predictive payoff distributions of different arms at time point *t*. It first defines a *safe set* of possible inputs 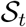 that are likely to provide outputs above the threshold, and then further separates the safe set into a set of *maximizers* (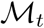, inputs that are likely to provide the maximum output) and *expanders* (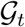, inputs that are likely to expand the safe set). Following Berkenkamp and Schoellig (2015), we define the upper and lower bounds of a confidence interval by adding the current expectation of reward 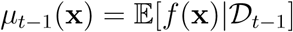 and the estimated uncertainty 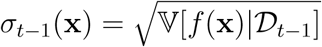:

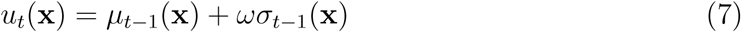

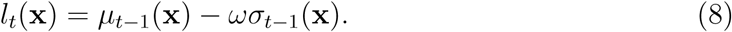

The parameter *ω* determines the width of the confidence bound, and we set it to *ω* = 3 to assure high safety for the rational safe optimization algorithm^3^. Using these bounds, we can define the safe set 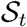 as all the input points in the set of available inputs 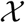 that are likely to lead to output values above the safe threshold *h*_min_,

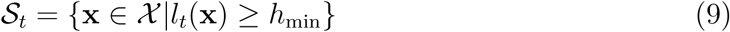

This means that points are considered to be safe if their lower confidence bound is above the provided threshold. This is intuitive as one would expect the output for these points to rarely fall below the threshold (around 99.9% of the times when setting *ω* = 3).

The set of potential maximizers 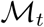 contains all safe inputs that are likely to obtain high output values; these are the safe inputs for which the upper confidence bound *u_t_* is above the best current lower bound (i.e. the highest lower bound of all input points):

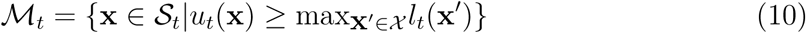

where **x**′ is the best revealed reward at time *t*. This means that maximizers are input points that are likely to be at least as good as the best overall outcome in a worst case scenario.

To find the set of expanders, we first define

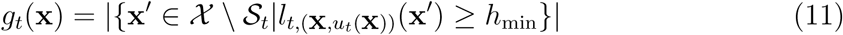

where *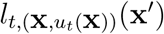* is the lower bound of **x**′ based on past data and a optimistic outcome for x which provides a new upper bound *u_t_* (**x**). Intuitively, this function is used to determine how many new inputs may potentially be added to the safe set after choosing input **x** and observing the output it provides. The function in (11) counts how many previously unsafe points can be classified as safe according to (9) assuming that *u_t_* (**x**) is measured when evaluating *f*(**x**). This function is positive only if the new data point has a non-negligible chance to expand the safe set. The set of potential expanders is then defined as

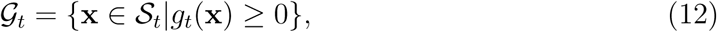

where *g_t_*(*x*) count the number of newly-introduced safe points (i.e. measures the cardinality of that set). The expander set is assessed by forward simulation and simply checks if the safe set is expected to be expanded (i.e., more points will be in the safe set after observing the expected outcome of the evaluated input) for a given input. Normally, the safe optimization algorithm operates by considering safe points that are either maximizers or expanders, and then choosing the point with the highest uncertainty about the expected outcome. However, here we use only a point’s membership in the three different sets in order to categorize participant behavior.

For categorizing participant decisions, we assess how much—if at all—choice behavior is guided by options being safe, maximizers, and/or expanders, as predicted by the safe optimization algorithm. More precisely, we check if membership of an input point within the three sets makes it more likely to be chosen. In order to assess if set membership is related to participant decisions, we use mixed-effects logistic regression to determine the extent to which factors of the safe optimization algorithm influence their choices. The dependent variable in this analysis is whether or not an option was chosen by a given participant on a given trial. The predictors are indicator variables for an option’s membership in the safe, the maximization, and the expander set. This analysis allows us to judge whether (i) a Gaussian process function learning model with parameters set to match the underlying task, combined with (ii) class membership specified by the safe optimization algorithm, can describe whether or not a participant chose an option on a given trial. This constitutes the first part of our analysis.

## Models of learning and decision making

In the second part of our analysis, we make out-of-sample predictions about individual choices, using a combination of different learning models with multiple decision strategies. We first contrast two different models of learning before describing the decision strategies. The *function learning* model learns about the underlying value function relating the spatial locations of options to their expected rewards. The *option learning* model does not learn about an underlying function, but rather learns about each input individually by associating inputs with previously generated rewards.

**Figure 1.**
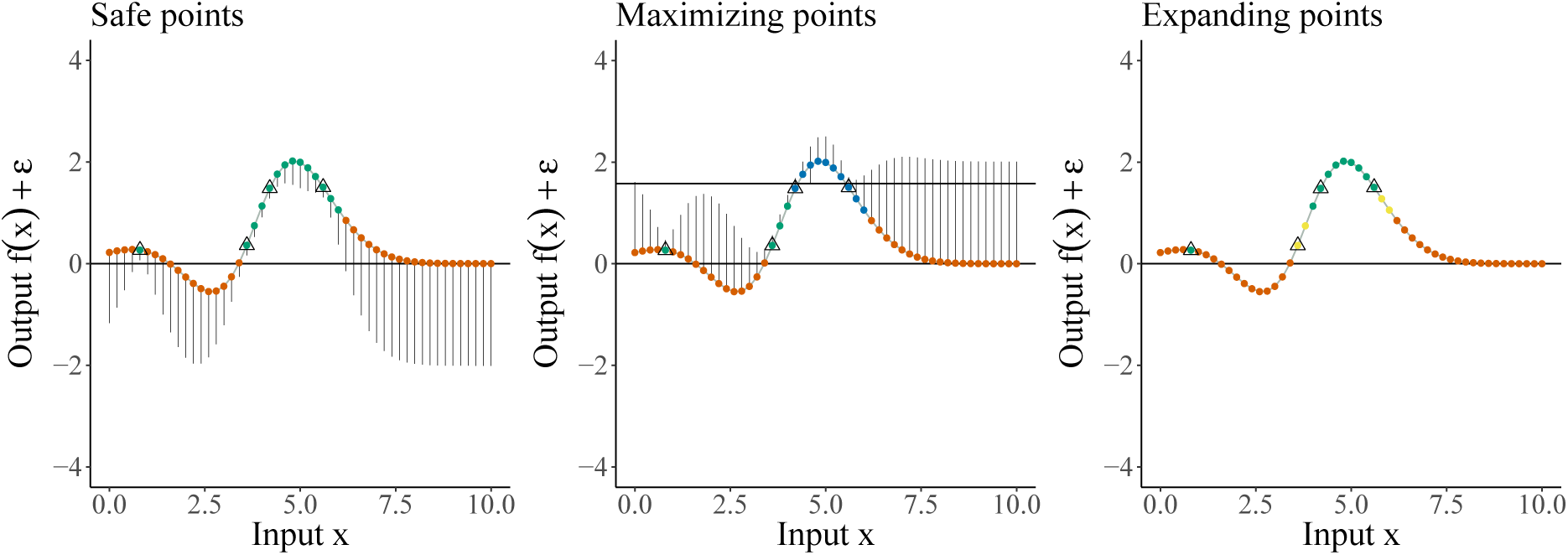
Example of sets estimated by safe optimization algorithm. Triangles indicate observations. Threshold was set to *h* = 0. **Left:** Identification of safe points (green), where the lower confidence bound (vertical lines) is higher than the threshold, and unsafe points (orange) where the lower confidence bound crosses the threshold. **Center:** Identification of maximizing points, which have an upper confidence bound (vertical lines) that is larger than the level (upper horizontal line) of the highest lower confidence bound across all points. Blue points represent maximizing points, green points represent safe points, and red points represent unsafe points. **Right:** Identification of expanding points, which are expected to increase the set of safe points when performing forward simulations. Yellow points represent expanding points, green points represent safe points, and red points represent unsafe points.

### Function learning model

For the function learning model, we use Gaussian process regression combined with a radial basis function kernel (Eq. 6). Using a radial basis function to model the extent of generalization across space is similar to Shepard et al. (1987)’s proposal of a universal law of generalization and has previously been implemented in a non-Bayesian model of function learning by Busemeyer, Byun, Delosh, and McDaniel (1997).

### Option learning model

The option learning model uses a simple mean tracking approach to learn the distribution of rewards of each input individually. We implement a version which assumes rewards are normally distributed with a known variance 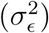 but unknown mean *θ_j_* where the prior distribution of the mean is again a normal distribution. This implies that the posterior distribution for each mean is also a normal distribution:

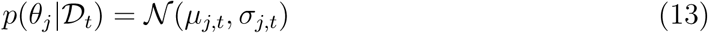

The mean *μ_j,t_* and variance 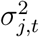 of the posterior distribution for option *j* are only updated when that option is selected at trial *t*:

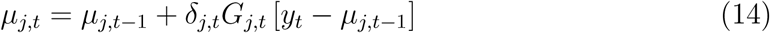

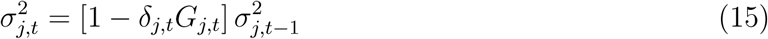

where *δ_j,t_* =1 if option *j* was chosen on trial *t*, and 0 otherwise. Intuitively, the estimated mean of the chosen option *μ_j,t_* is updated based on the difference between the observed value *y_t_* and the expected mean *μ_j,t−_*_1_, multiplied by *G_j,t_*. At the same time, the estimated variance 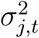 is reduced by a factor of 1 − *G_j,t_*, where *G_j,t_* is defined as:

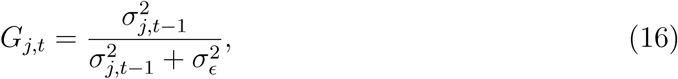

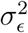 is the error variance, which is estimated as a free parameter per round. We set the prior mean to the median value of the payoffs and the prior variance 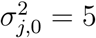

This model does not generalize over unseen arms at all, but rather only learns locally about the distribution of rewards for each option separately (Wu et al., 2018). It can also be considered as a special case of the function learning model as the assumed correlation between points goes to zero. We use this model as a benchmark for comparisons in our cross-validation procedure. If this model predicts participant behavior well, this means that participants do not generalize using the spatial structure of the environment, but rather learn about each option independently, as is the case in a traditional multi-armed bandit.

### Decision strategies

The learning models produce predictions about the distribution of rewards for each option in the search space, whereby we use a decision strategy to determine useful actions and predict choices. We compare four different strategies, two of which are designed for safe search and two of which are designed for risky search. Each strategy is based on an acquisition function, which determines a utility value for each option, with a softmax choice used to make probabilistic predictions about choice behavior.

**Decision strategies for safe tasks**. *Upper confidence bound sampling* directly trades off between the expected rewards and uncertainty. Given the posterior mean *μ_t_*_−1_(**x**) and its attached standard error *σ_t_*_−1_(**x**), we calculate the acquisition function of the upper confidence bound as

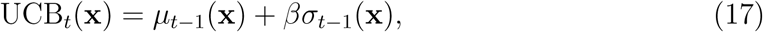

where the exploration factor *β* determines how much reduction of uncertainty is valued (relative to exploiting known high-value options) and is estimated as a free parameter. This sampling strategy has recently been found to describe human behavior well across different function exploration-exploitation tasks without additional constraints (Schulz et al., 2017; Wu et al., 2018). Additionally, it has known performance guarantees in function optimization scenarios (Srinivas, Krause, Kakade, & Seeger, 2012). We use it as a candidate for unconstrained function optimization tasks.

The *probability of improvement* (POI) strategy evaluates an option based on how likely it will be better than the best outcome (**x**^+^) observed so far:

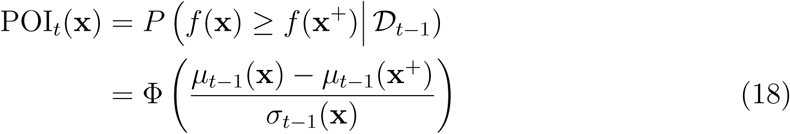

where Φ(·) is the normal CDF. This rule calculates the probability for each option to lead to an outcome higher than the option that has currently been observed (Kushner, 1964) and has recently been used in experiments involving multi-attribute choices (Gershman, Malmaud, & Tenenbaum, 2017).

**Decision strategies for risky tasks**. To define possible search strategies for risky situations, we consider two modifications of the decision strategies defined above. The *probability of being safe* (POS) is similar to the POI strategy, but assess the probability that a candidate input provides a reward above the safe threshold.

Formally, if the threshold is *h*_min_, POS is defined as:

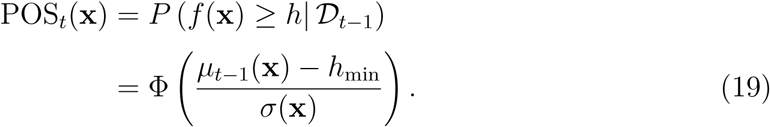

Instead of sampling by the probability to improve upon the best seen point so far, this sampling strategy only cares about maximizing the probability of being safe (i.e., sampling above the threshold). This strategy is very risk averse and frequently prefers known options over exploratory choices.

Instead of valuing uncertainty positively (as is the case with UCB), the *lower confidence bound algorithm* (LCB) tries to avoid highly uncertain options:

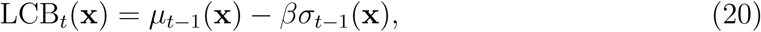

Because inputs with high uncertainty can also lead to possibly bad outcomes, this sampling strategy can be seen as highly risk-averse but possibly not very adaptive approach to risky environments. The difference between UCB and LCB also corresponds to differences observed in risk-sensitive reinforcement learning when outcomes are either positive or negative (Niv, Edlund, Dayan, & O’Doherty, 2012). A related sampling strategy that can account for the possibility of a negative uncertainty bonus (i.e., valuing uncertainty as bad) in the domain of losses has been considered by Krueger, Wilson, and Cohen (2017) before.

### Estimation and model comparison

For model fitting and evaluation, we use a cross-validation procedure in which we fit the model using maximum likelihood estimation on a subset of the data, and then use the estimated parameters to make out-of-sample predictions on the remaining data. For each model, we use a softmax function to transform each model’s criterion into a probability distribution over options:

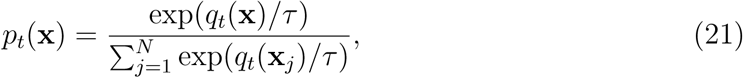

where *q_t_*(**x**) is the value of an option **x** according to each model, and *τ* is the temperature parameter (i.e., lower values of *τ* indicate more precise predictions).

For the function learning model, we estimate λ (length-scale), for the option learning model 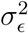 (error variance), and for the upper and lower confidence bound sampling strategies *β* (exploration bonus for UCB, safety bonus for LCB). Additionally, all models include *τ* as a free parameter. We fit all models separately for each participant by cross-validated maximum likelihood estimation, using a differential evolution algorithm (Mullen, Ardia, Gil, Windover, & Cline, 2009). Parameter estimates are constrained to positive values in the range [exp(−5), exp(5)].

Cross-validation is performed for the safe and risky function exploration-exploitation objectives separately. Within all rounds, we use leave-one-block-out cross-validation to iteratively form a training set by leaving out a single round, computing a maximum likelihood estimate on the training set, and then generating out-of-sample predictions on the remaining round. This is repeated for all combinations of training and test sets, and for every participant individually. The prediction error (computed as log loss) is summed up over all trials, and is reported as average *predictive accuracy*, using a pseudo-*R*^2^ measure that compares the total log loss prediction error for each model to that of a random model

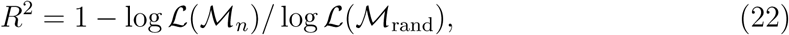

where log 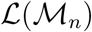 is the log loss (negative log likelihood) of model *n* and log 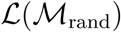 the log loss of a random model (which chooses options with equal probability). Intuitively, a *R*^2^ = 0 corresponds to prediction accuracy equivalent to chance, while *R*^2^ = 1 corresponds to a perfect prediction accuracy.

## Experiment 1: Univariate inputs

The first experiment required participants to maximize unknown univariate functions *f*: *x* ↦ *y* by choosing discretized input values *x* ∈ (0, 0.5, 1,…, 10). This scenario is similar to a multi-armed bandit task (with *n* = 21 arms) in which all arms are ordered horizontally and where the outputs of the arms are correlated as a function of their distance. Additionally, we introduced the constraint that participants should avoid choosing options with a reward below the horizontal red line, or else forfeit the remaining trials in the round.

### Participants

61 participants (36 female) with an average age of 32.95 (SD = 8.02) were recruited via Amazon Mechanical Turk and received $1.00 USD for their participation and a bonus of up to $1.00 in proportion to their overall score. The experiment took on average 12 minutes to complete.

### Procedure

Participants were told they had to maximize outputs of an unknown function, while at the same time trying to avoid obtaining outputs below a given red line. After reading the instructions and performing an example round, they had to correctly answer 4 comprehension questions to check if they understood the instructions. There were 9 rounds in total and each round contained (at most) 10 trials. At the start of each round, participants were shown the output of a single preselected input value, which was randomly sampled from all inputs with outputs above the threshold *h*_min_.

On each trial, participants were asked to choose an input with an output above the red line (i.e., a “safe” option), and told that choosing an input below the line would end the current round, forfeiting potential additional rewards they could have earned by finishing the round. On each trial *t =* 1,*…*, 10 in a round, they could choose an input value *x* ∈ (0, 0.5, 1,*…*, 10} to observe (and acquire) a reward *y* = *f*(*x*) + ∊ with noise term 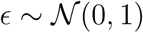. The underlying functions were sampled from a GP prior with a radial basis function kernel (length-scale λ=1). Participants were told that the objective was to maximize the sum of the obtained rewards over all trials in a round (i.e., score), which corresponded to a bonus of *score* × $0.09.

Before the first trial, an initial safe point above the threshold was sampled at random and provided to participants. A screenshot is shown in Figure 2. Rewards were scaled to be between 0 and 10 but such that the underlying maximum was never actually 10 in order to make the maximum not easily guessable. This was done by sampling a random number between 9 and 10 and using this number as the overall maximum for rescaling.

**Figure 2.**
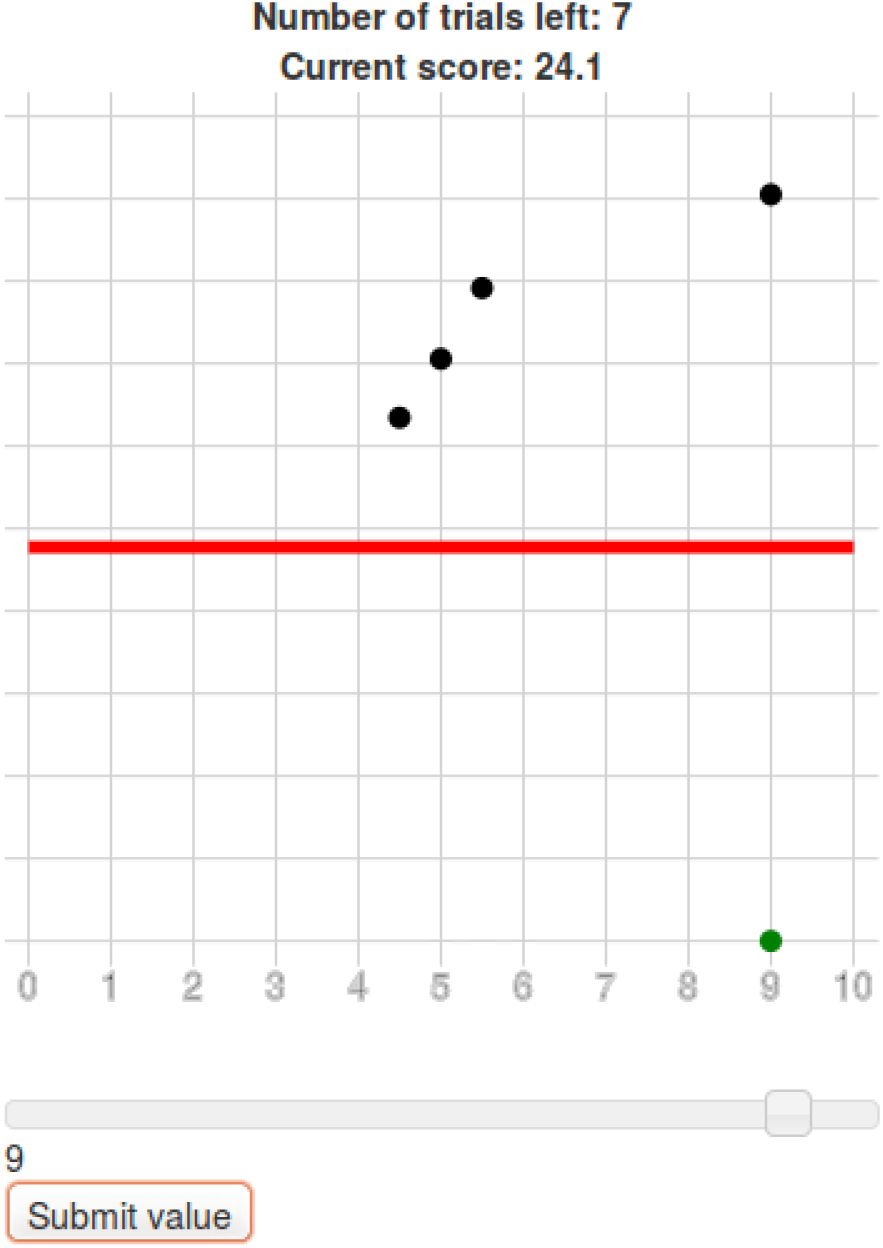
Screenshot of Experiment 1. The red line marks the safe threshold (outcomes below this threshold immediately ended the round). Dots above the red line show observed outputs. Inputs were chosen by moving the slider and selected by clicking “submit value”, with the green dot indicating the observed reward.

In order to see if different levels of risk influence participant learning and sampling behavior, we manipulated the risk of obtaining outcomes below the red line as a between-group factor, resulting into 6 groups for which the probability of sampling below the line was set to *p* = [0.55, 0.6, 0.65, · · ·, 0.8]. This means that, unknown to participants and before the start of each round, the red line was placed such that proportion *p* of the input points would produce an output lower than the red line, corresponding to the different risk conditions.

### Behavioral results

Figure 3 shows the results of Experiment 1. In general, participants performed better than chance (mean score = 6.21, *t*(61) = 12.32, *p <* .001, *d* = 1.57) and improved over time (mean correlation between trials and score *r* = .2, *t*(60) = 6.87, *p <* .001, *d* = 0.88). In addition, the average number of trials per block statistically exceeded what would be expected if participants chose completely at random (*t*(60) = 22.69, *p <* .01, *d* = 2.9), indicating that participants were successful at finding reward and avoiding risky options. Participants assigned to different risk conditions did not perform significantly different from each other (correlation between mean score and risk level: *r* = .06, *t*(59) = 0.48, *p >* .6). Participants also showed localized sampling behavior, choosing inputs more locally than a random sampling model (*t*(60) = −22.1, *p <* .001, *d* = −2.83), although participants in higher risk conditions did not choose more locally (correlation between risk level and average distance of consecutive inputs: *r* = −.08, *t*(59) = −0.64, *p* = .52). Therefore, participants learned within the task but were seemingly uninfluenced by the riskiness of the threshold, perhaps because they stayed almost exclusively to safe points in all of the threshold conditions.

**Figure 3.**
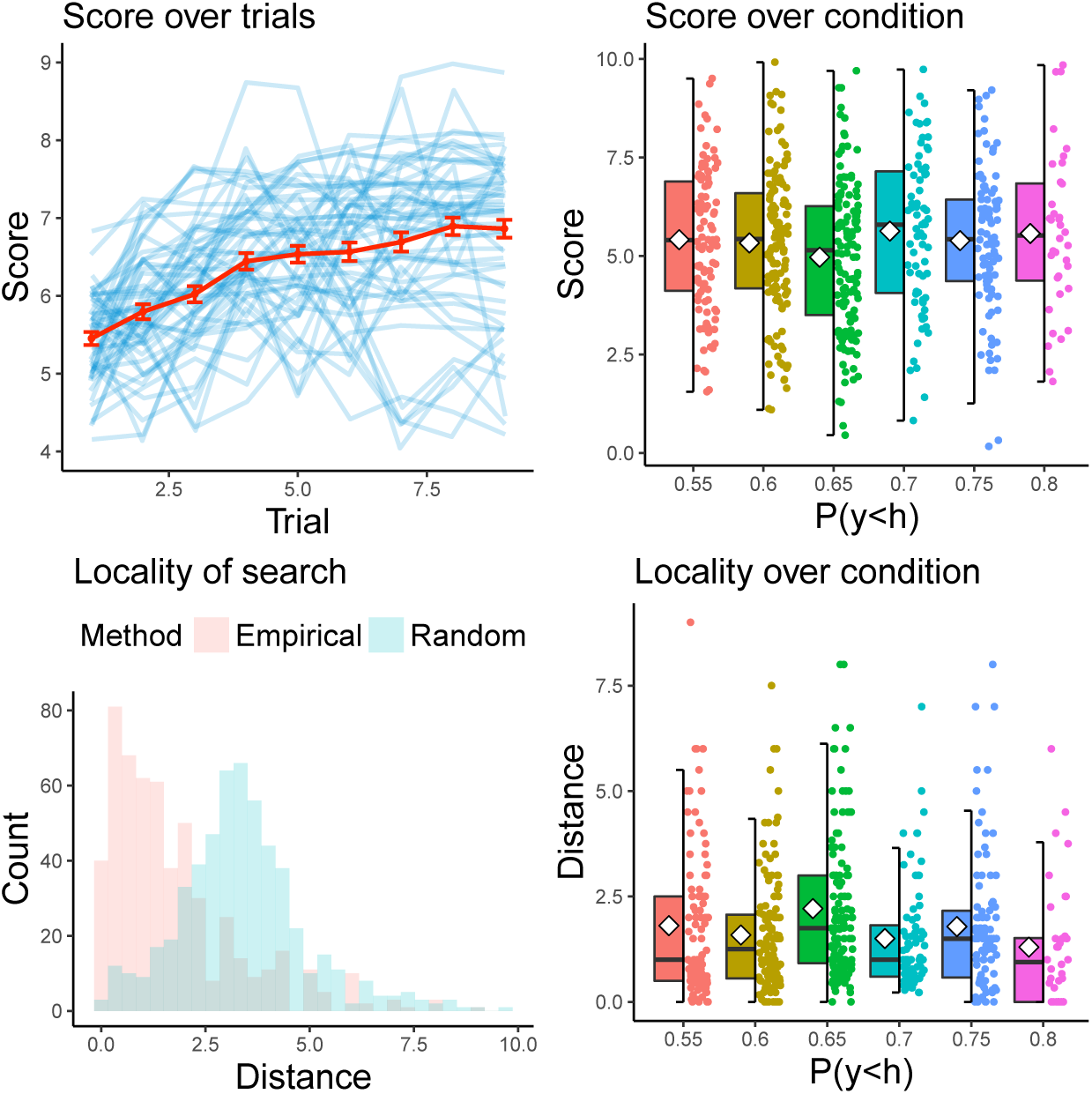
Results of Experiment 1. The upper left panel shows participants’ scores over trials, including the average score (red) and standard errors in error bars. The upper right panel shows a box plot of participants’ scores for the different risk conditions including raw data points and group means (white diamonds). The lower left panel shows the locality of chosen inputs as compared to a random sampler. The lower right panel shows a box plot of the locality of chosen inputs by different riskiness-conditions including raw data points and group means (white diamonds).

### Categorization of decisions

We used mixed-effects logistic regression analysis to determine the factors influencing participant decisions. The dependent variable was whether each input was chosen or not on each trial for each participant. As predictors, we used indicator variables for an input’s membership in the safe, maximization, and expander sets. Results indicated that the most plausible model was one that contains all variables as fixed effects and a participant-specific random intercept, indicating that participants were influenced by set membership in an overall similar fashion. The coefficients of the fixed effects are presented in Table 1 below.

**Table 1.** Results of the mixed effects logistic regression for Experiment 1.

|  | Estimate | S.E. | z value | Pr(>\|z\|) |
| --- | --- | --- | --- | --- |
| Intercept | -3.71 | 0.12 | -29.75 | 0.00 |
| Maximizer | 0.82 | 0.23 | 3.50 | 0.00 |
| Expander | -0.44 | 0.20 | -2.20 | 0.03 |
| Safe | 0.85 | 0.24 | 3.46 | 0.00 |

Comparing the magnitude of the slopes of the predictors, we can conclude that while all of the sets relate to participant behavior, participants were mostly influenced by whether or not a point was safe (Estimate = 0.85) or a maximizer (Estimate = 0.82). Being within the set of possible expanders was negatively related to whether or not participants would choose a given point (Estimate = −0.44). This shows that participant behavior corresponded, at least to some extent, to the predictions generated by the rational model. However, they seemed to focus more on staying safe and maximizing locally rather than expanding the safe set. We next assessed if we could predict trial-by-trial decision behavior with more process-level modeling.

### Trial-by-trial models

Assessing the trial-by-trial modeling results, we found that the overall best performing models were the Gaussian process model with the probability of being safe decision strategy as well as the option learning model with the same decision strategy. Comparing the learning models aggregated over decision strategies, we found that the option learning model outperformed the Gaussian Process learning model (*t*(60) − 2.34, *p <* .05, *d* = 0.30). This mirrors the behavioral finding that participants explored in a local manner, thus seemingly not generalizing much over different inputs and observed outcomes. Comparing the decision strategies aggregated over learning strategies, we found that the probability of being safe strategy predicted participant behavior better than any of the other strategies, no matter whether it was combined with the option learning (*t*(60) = 5.72, *p <* 0.001, *d* = 0.73) or the Gaussian process function learning model (*t*(60) = 6.28, *p <* 0.001, *d* = 0.80). Finally, there was no significant difference between the Gaussian process function learning and the option learning, when both were combined with the probability of being safe sampling strategy (*t*(60) = −0.71, *p* = 0.48, *d* = 0.09).

We extracted the median parameter estimates of the Gaussian process learning model combined with the probability of being safe sampling strategy for each participant to check if they meaningfully tracked behavioral differences in the task (Figure 5). Overall, the model’s predictions were relatively precise as indicated by low estimates of the softmax temperature parameter *τ* (median estimate: 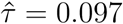). Furthermore, the length-scale parameter of the Gaussian process indicated that participants seemed to somewhat generalize over different arms (median estimate 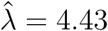; compared to the ground truth of λ =1). Interestingly, people who generalized more performed worse overall (*r* = −.36, *t*(59) = −2.99, *p <* .004). This is most likely the result of the dual objectives participants were facing, which required them to generalize beyond encountered examples but to also sample safe options, which frequently required sampling rather locally.

**Figure 4.**
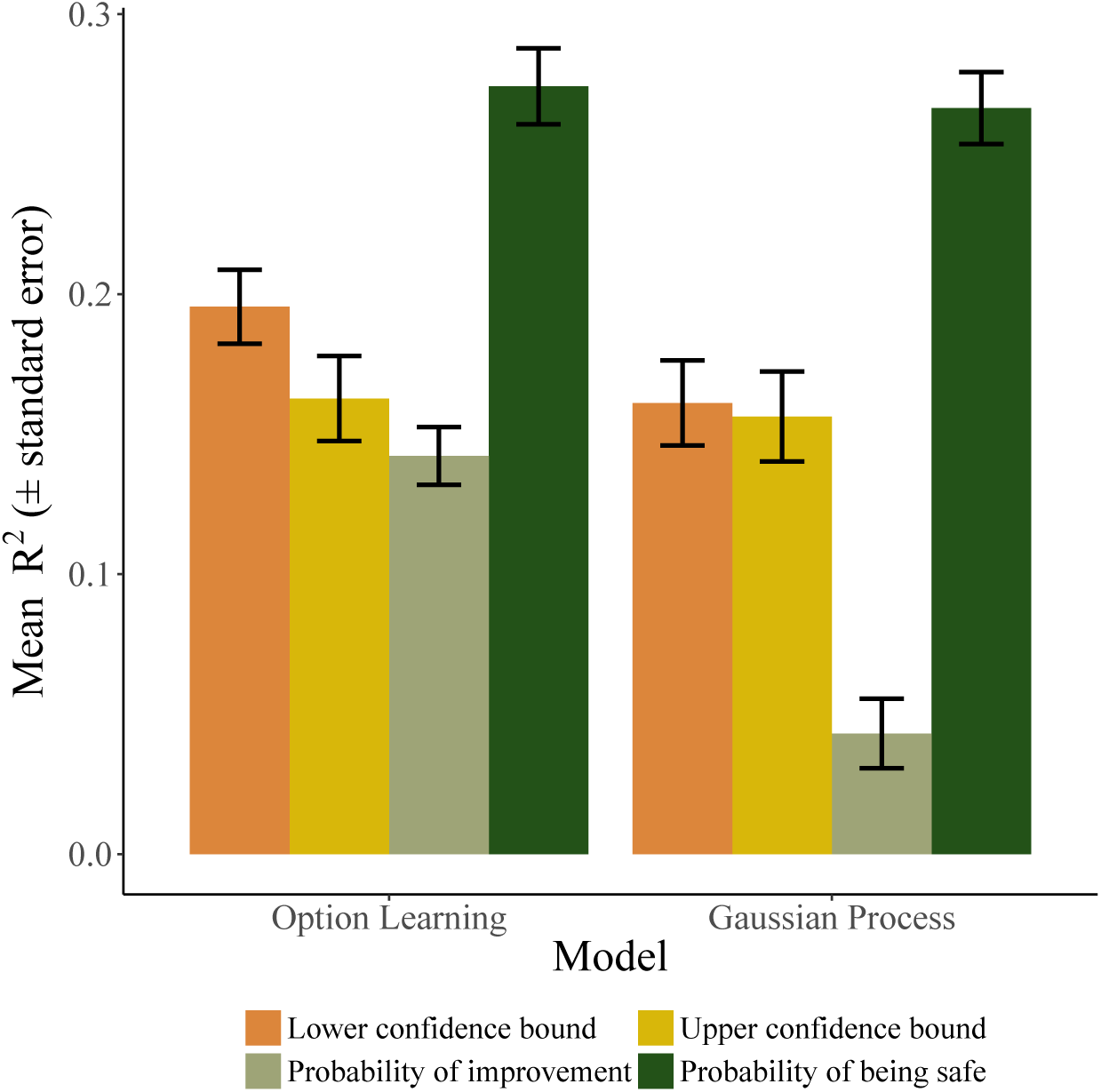
Results of the trial-by-trial learning models in Experiment 1.

**Figure 5.**
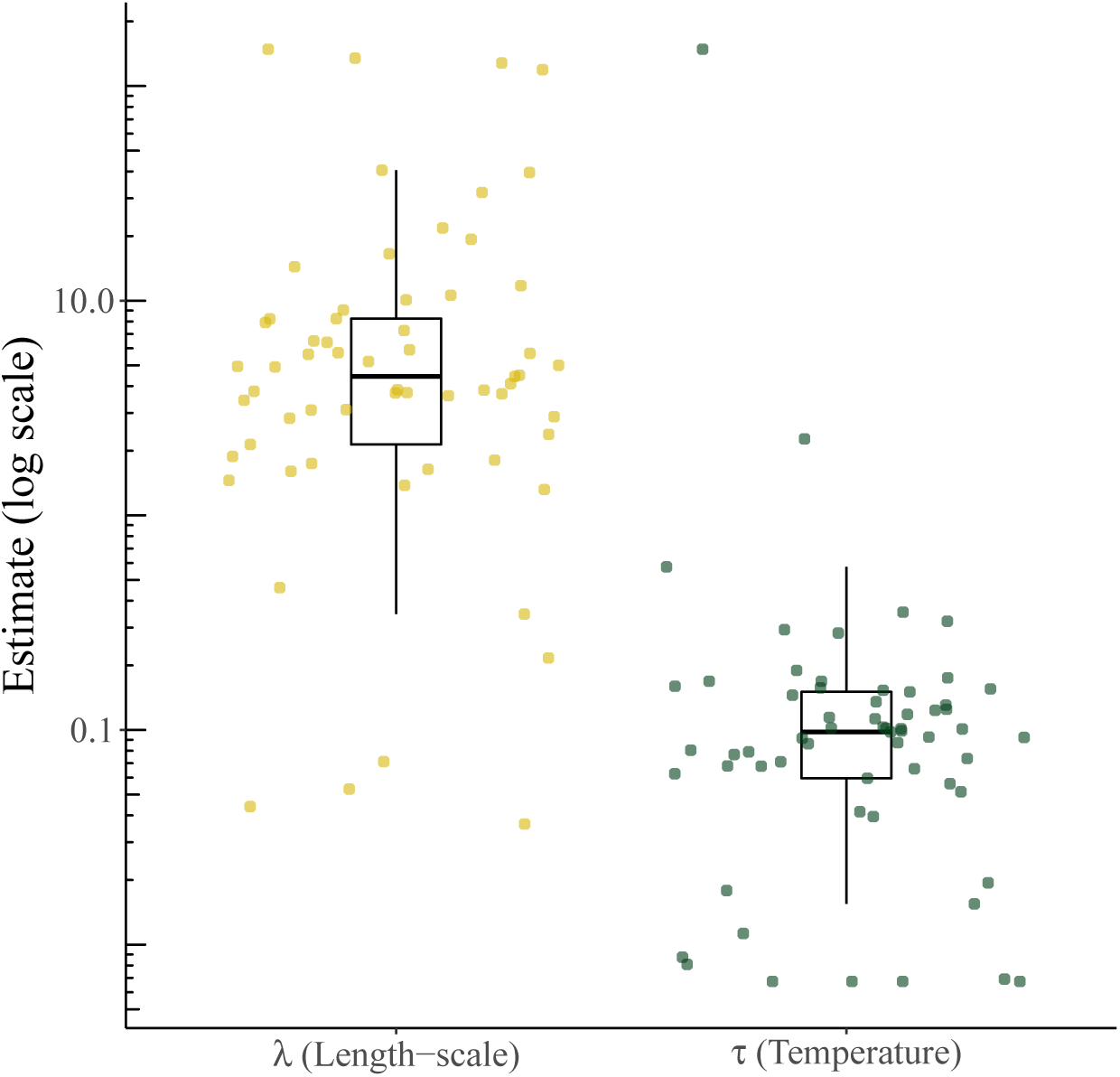
Parameter estimates of the Gaussian process and probability of being safe (GP-POS) model in Experiment 1.

Lastly, we used the individual participant parameters estimates of the Gaussian process function learning model paired with the probability of being safe sampling strategy (henceforth GP-POS) to simulate data within the task. This means that we let specified the model using participant estimates for both *λ* and *τ*, and simulated performance in the exact same task as participants, for the exact same number of rounds, trying to optimize the same underlying function. The results of this posterior model simulation allow us to assess the extent to which the empirical results can be reproduced by the GP-POS model. The results of this simulation are shown in Figure 6.

**Figure 6.**
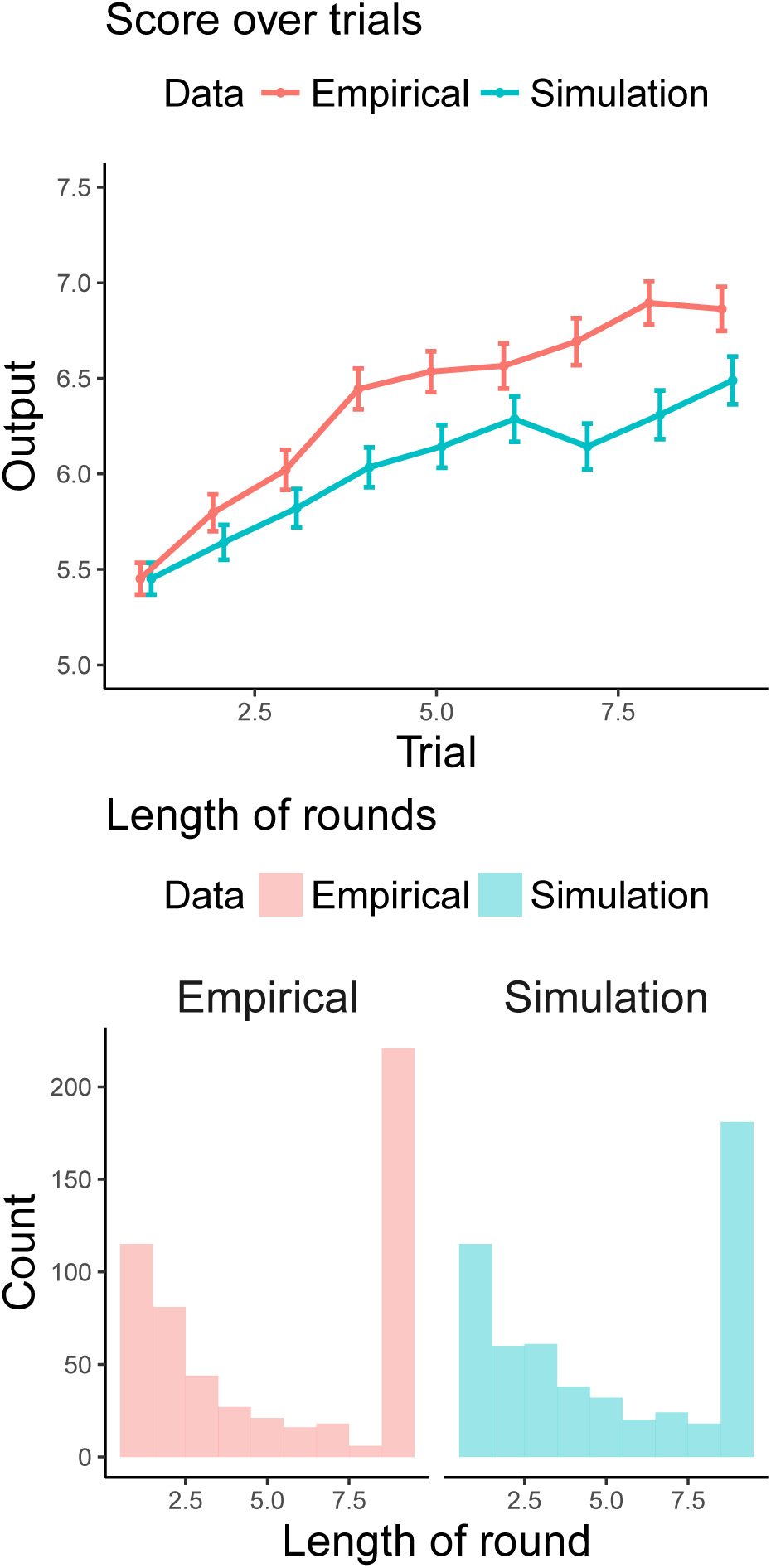
Posterior model checks of GP-POS model for Experiment 1. Upper panel shows the mean trajectories of scores over trials for both human participants and the GP-POS model when performing the exact same task. Lower panel shows the histogram of lengths per round, i.e. how long both participants and the model managed to sample without sampling below the threshold.

Even though the GP-POS model produces a similar trajectory of mean rewards as participants, its average score per trial is somewhat below that of participants. Looking at the distribution of average trial length per round, we can see that while the two distributions are very similar overall, participants managed to more often successfully complete the full number of rounds (i.e., without sampling below the threshold) than the simulations. While this could indicate that participants were even more risk averse than the best fitting GP-POS models, the finding that they also outperformed the GP-POS model shows that this did not negatively affect their performance.

## Discussion

Within a first experiment assessing behavior in a univariate variant of the risky spatially correlated multi-armed bandit, we found that participants managed to successfully learn within this tasks, improved their scores over trials, and performed better than expected by chance. Moreover, participants tended to select input points which were classified as safe or maximizing points by a rational safe optimization algorithm. Using leave-one-block-out cross-validation, we found that participant behavior was best predicted by a probability of staying safe (POS) decision strategy is primarily concerned with sampling points above the provided threshold. Overall, participants did not generalize far beyond the already observed input points, resulting in roughly equal performance of both the option learning and the Gaussian process regression model. There seemed relatively little effect of the level of riskiness in the task (manipulated by the level of the safe threshold). Perhaps participants start behaving equally risk averse once a threshold is introduced. Overall, participants performed slightly better and chose inputs in a more risk averse manner than predicted by the GP-POS model (parameterized by participant’s estimates), although the model produced human-like behavior overall. To further discern whether there is continuous adaptation or whether the introduction of any threshold leads to similar behavioral outcomes we will apply a within-subject manipulation of the level of riskiness in Experiment 2.

## Experiment 2: Bivariate inputs

In the second experiment, participants were asked to maximize an unknown bivariate function, which was represented by a two-dimensional grid world (Fig. 7). Moreover, we introduced a standard, risk-free condition as a within-subjects factor to see if participants can switch between the two different modes or riskiness.

**Figure 7.**
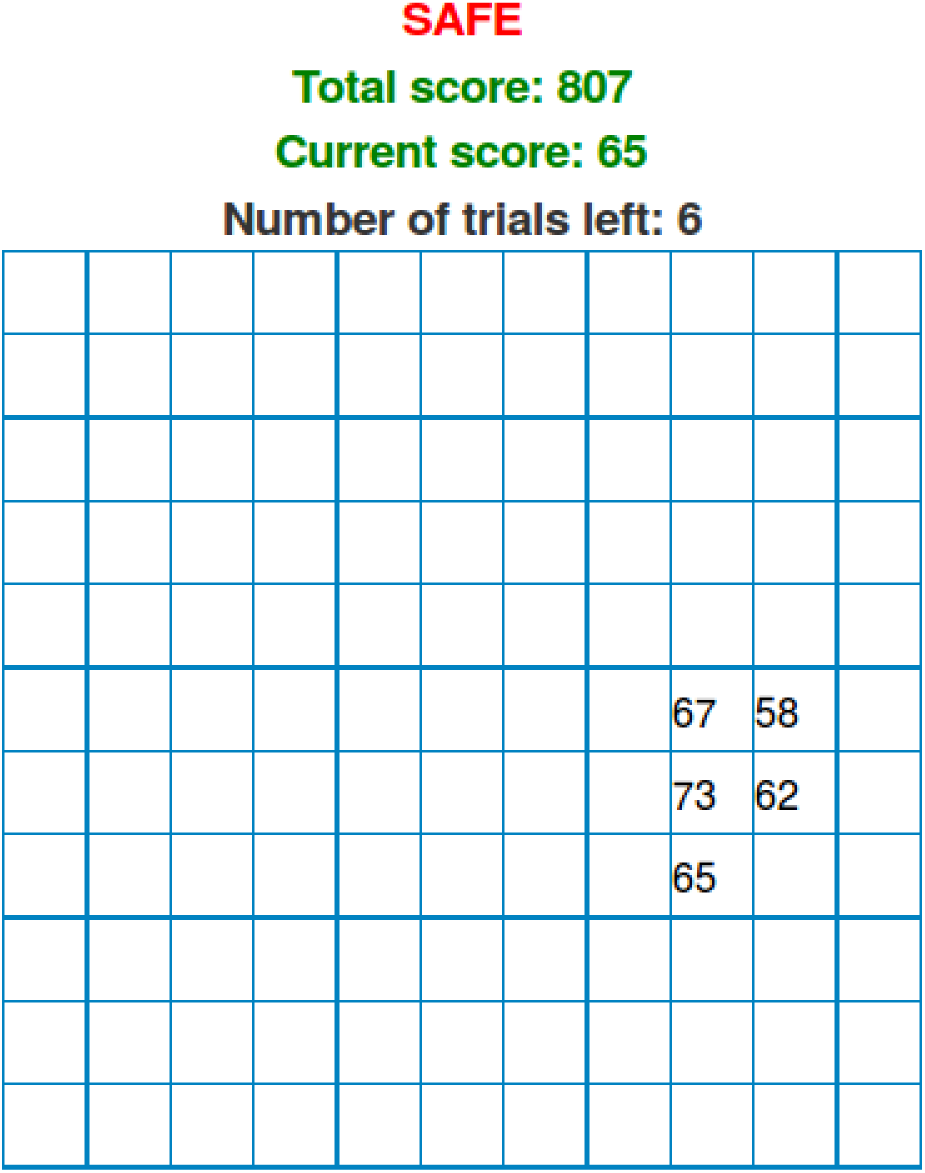
Screenshot of Experiment 2. Inputs were arranged in an 11 by 11 grid. Participants chose inputs by clicking on the corresponding tile, trying to choose inputs which produce high rewards. The “SAFE” condition indicates that they do not have to worry about obtaining inputs above the safe threshold of 50. Rounds at which they had to obtain outputs above 50 were marked as “RISKY”.

### Participants

62 participants (37 male), with an average age of 31.77 years (SD = 8.97) were recruited via Amazon Mechanical Turk and received $1.00 USD for their participation and a performance-dependent bonus of up to $1.00 USD. The average completion time of the whole experiment was 11 minutes.

### Procedure

We created functions *f*: **x** ↦ *y* with **x** = (*x*_1_,*x*_2_)^⊤^, defined over the grid *x*_1_*, x*_2_ ∈ [0, 0.1,…, 1] resulting in a 11 × 11 grid, with *y* = *f* (**x**) + *∊* with 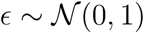. As in Experiment 1, a function *f* was sampled independently from a GP with an RBF kernel (length-scale λ = 2) on each round. The output values *y* varied between 0 and 100 and one initial input point with an output above the threshold of 50 was chosen at random and provided at the start of each round. We varied the level of risk within-participants: out of the total of 10 rounds there were 5 which were “safe” (i.e., unconstrained maximization tasks without a threshold) and 5 which were labeled as “risky” (i.e., constrained maximization tasks where obtaining an output below 50 caused the round to terminate immediately, forfeiting any remaining trials in the round). The rounds were presented in a counter balanced order. Participants were paid a basic fee of $1 and an additional bonus of $0.01 for every 10 points they earned overall.

### Behavioral results

Figure 8 shows the results of Experiment 2. Participants performed better than chance overall (*t*(61) = 15.48, *p <* .001, *d* = 1.97). On average, participants did not increase their scores significantly over trials (mean correlation: *r* = .04, *t*(61) = −1.34, *p >* .1, *d* = 0.16). However, looking separately at the riskiness conditions showed that while this was true for the safe conditions (mean correlation: *r* = .02, *t*(61) = 0.99, *p >* .3, *d* = 0.13), participants did get significantly better over trials within the risky condition (mean correlation: *r* = .10, *t*(61) = 2.23, *p <* .05, *d* = 0.28). Surprisingly, participants scored higher in risky rounds compared to safe rounds (*t*(61) = 9.78, *p <* .001, *d* = 1.24). This seems to be driven by a tendency towards greater exploration of the whole input space in the safe rounds which also explains why the average of their first sampled output considerably drops from the average revealed value (see Figure 8). In risky rounds, participants avoided scoring below the threshold for longer than expected by chance (*t*(61) = 8.06, *p <* .001, *d* = 1.02).

**Figure 8.**
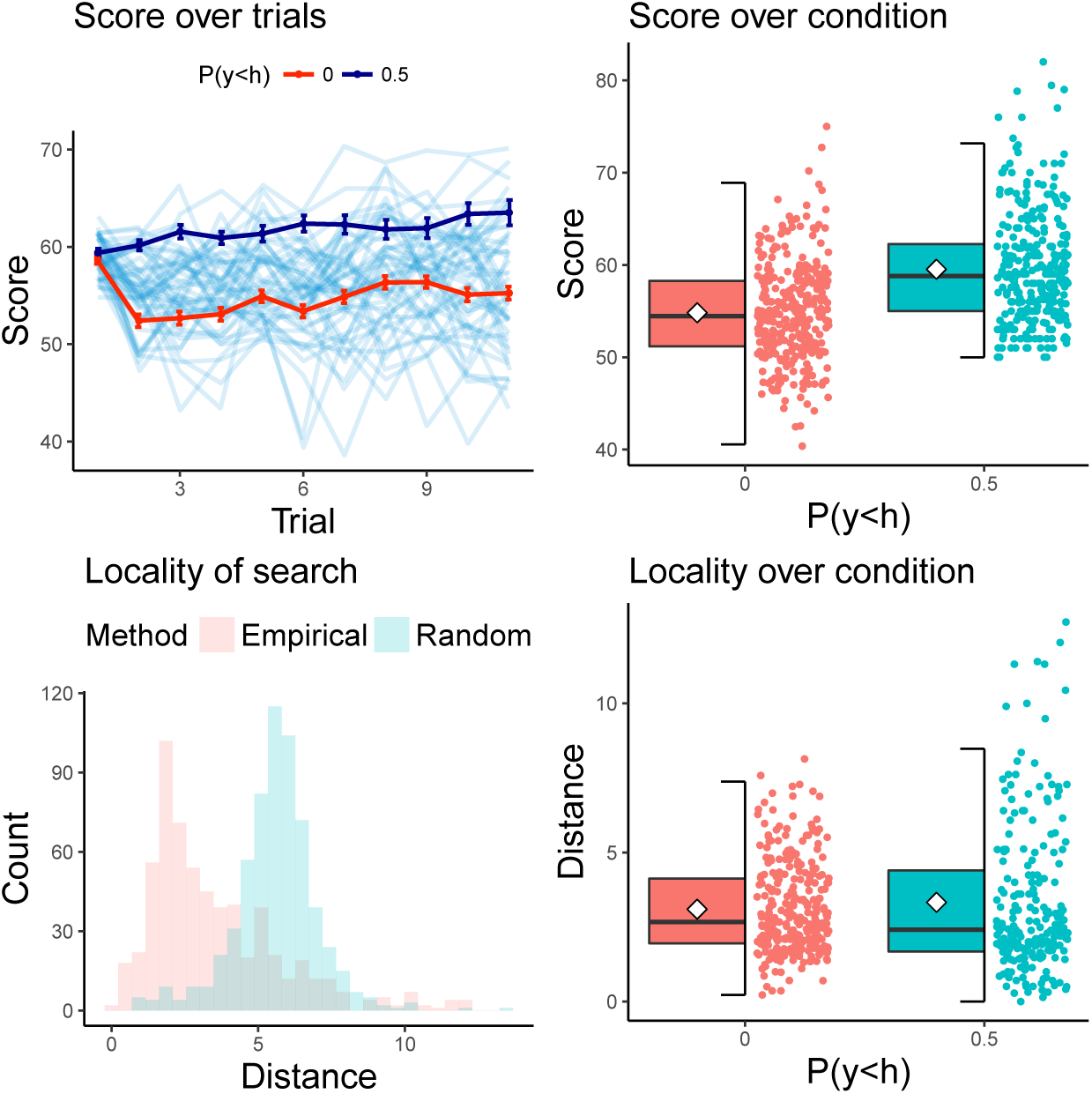
Results of Experiment 2. The upper left panel shows participants’ scores over trials, including the average score (red lines, blue line shows mean of risky condition, red line of safe rounds) and standard errors in error bars. The upper right panel shows a box plot of participants’ scores for the risky and safe condition including raw data points and group means (white diamonds). The lower left panel shows the locality of chosen inputs as compared to a random sampler. The lower right panel shows box plot the locality of chosen inputs for the safe and risky rounds.

Participants again explored more locally than expected by chance (*t*(61) = 18.43, *p <* 0.001, *d* = 2.34), but did not explore more locally during risky as compared to safe rounds (*t*(61) = −0.31, *p >* 0.7, *d* = −0.06). Thus, participants might have sampled further away from the first point at the beginning, but later on again sampled rather locally.

### Categorization of decisions

We again fit a mixed-effects logistic regression analysis to participants’ choices and found that the best possible model contained a random intercept over participants as well as an interaction term between the effect of the safe sets and the current riskiness condition (Table 2). As expected, the effect of the safe set was again the strongest overall (Estimate = 1.85). As before, participants also chose points that were classified as maximizers more frequently (Estimate = 1.16). Additionally, being within the expansion set also deemed points to be significantly more frequently chosen this time, although this effect turned out to be relatively small (Estimate = 0.13). The interaction effect between the riskiness condition and the safe sets indicate that participants are less likely to sample from within the safe sets in the conditions without additional risks (Estimate = −0.61). This is intuitive as they are not required to focus on sampling above 50 in this condition.

**Table 2.** Fixed effects of the mixed-effects logistic regression in Experiment 2.

| | Estimate | S.E. | z value | $\Pr(>\vert z\vert )$ |
| --- | --- | --- | --- | --- |
| Intercept | -5.17 | 0.02 | -271.95 | 0.00 |
| Maximizer | 1.16 | 0.11 | 10.58 | 0.00 |
| Expander | 0.13 | 0.06 | 2.24 | 0.02 |
| Safe | 1.85 | 0.11 | 16.75 | 0.00 |
| Safe × Condition | -0.61 | 0.06 | -10.62 | 0.00 |

### Trial-by-trial modeling

Model fits for the two riskiness conditions are shown in Figure 9. We can see that the predictions of the models were generally better for the risky condition than the safe conditions (*t*(61) = 9.69, *p <* .001, *d* = 1.23). Only analyzing the safe condition, we found that the Gaussian process regression model led to better predictions than the option learning model (*t*(61) = 4.89, *p <* .001, *d* = 0.62). There were no differences between the different decision strategies when paired with the Gaussian process model for the safe conditions (all *p >* 0.05). Looking at the results for the risky condition, the Gaussian process regression again predicted behavior better than the option learning model (*t*(61) = 4.53, *p <* .001, *d* = 0.58). Importantly, the probability of being safe sampling strategy led to significantly better prediction than any other sampling strategy (*t*(61) = 3.37, *p <* .01, *d* = 0.43). Therefore, participants seem to adapt their sampling strategy to the risky constraints of the task.

**Figure 9.**
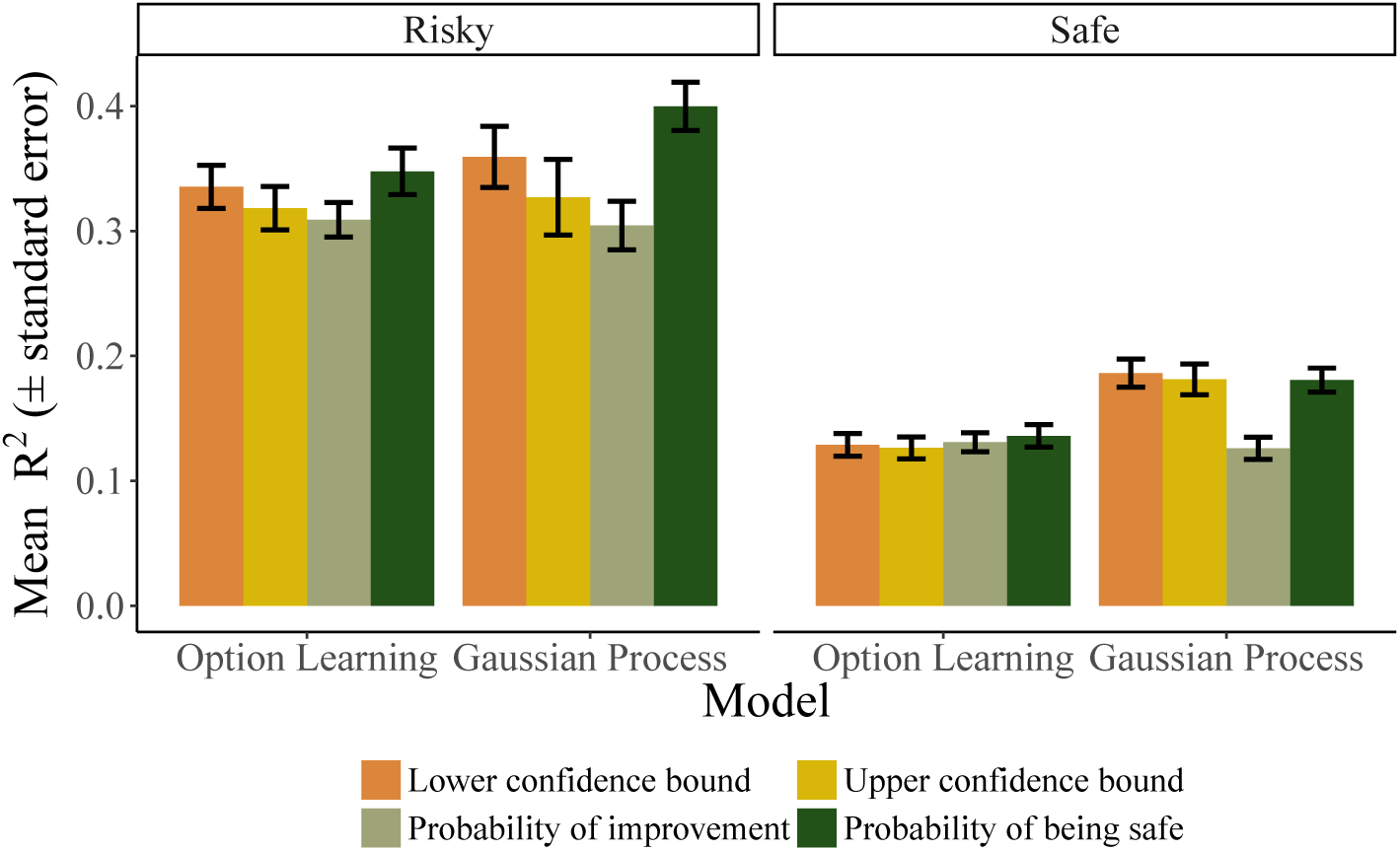
Trial-by-trial modeling results for Experiment 2.

Figure 10 shows the median parameter estimates for each participant for the GP-POS model(Gaussian process learning paired with the probability of being safe decision strategy) for both the risky and safe conditions. Whereas the estimates of the softmax temperature parameter differed between the two conditions (*t*(61) = −4.13 *p <* .001, *d* = 0.52) indicating more precise predictions for the risky (median estimate 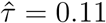) than for the safe condition (median estimate 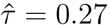), estimates of the length-scale λ did not differ significantly between conditions (*t*(61) = −0.66, *p* = .51, *d* = 0.08). Instead, median estimates of λ per participant correlated significantly between conditions (*r* = .36 *t*(60) = 2.97, *p <* .01), indicating that participants approached both conditions with a similar tendency towards generalization.

**Figure 10.**
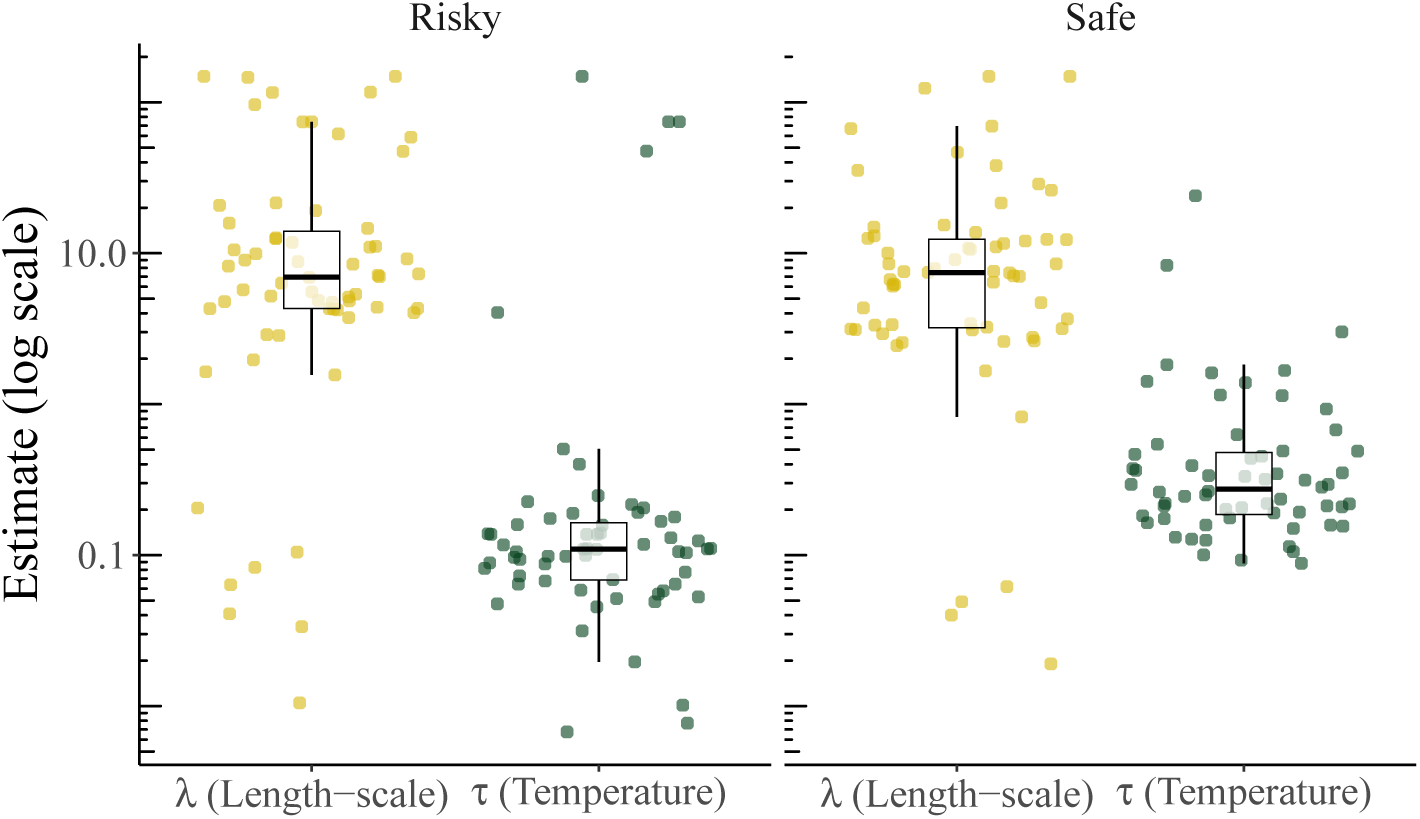
Parameter estimates of the GP-POS model in Experiment 2.

As in Experiment 1, we performed a posterior model check by parameterizing the GP-POS model with the participant-wise parameter estimates and let the model perform the exact same task as participants. Results of this simulation are shown in Figure 11. We can see that, in the risky condition, the average score of the GP-POS model drops on the second trial but afterwards increases more sharply than participants’ mean trajectory. From round 9 onwards, the model performs slightly better than participants. This indicates that the GP-POS model explores more extensively than participants did, incurring an initial hit to performance in order to reap later benefits. For the safe condition, the model corresponds almost perfectly with participants’ mean trajectories. This is expected as this result is primarily driven by a higher temperature parameter *τ*, leading to an increase in random exploration. The histogram of round length again indicates that participants’ behavior is more risk averse than that of the GP-POS model, as participants managed to play for the maximum number of trials more frequently than he GP-POS model.

**Figure 11.**
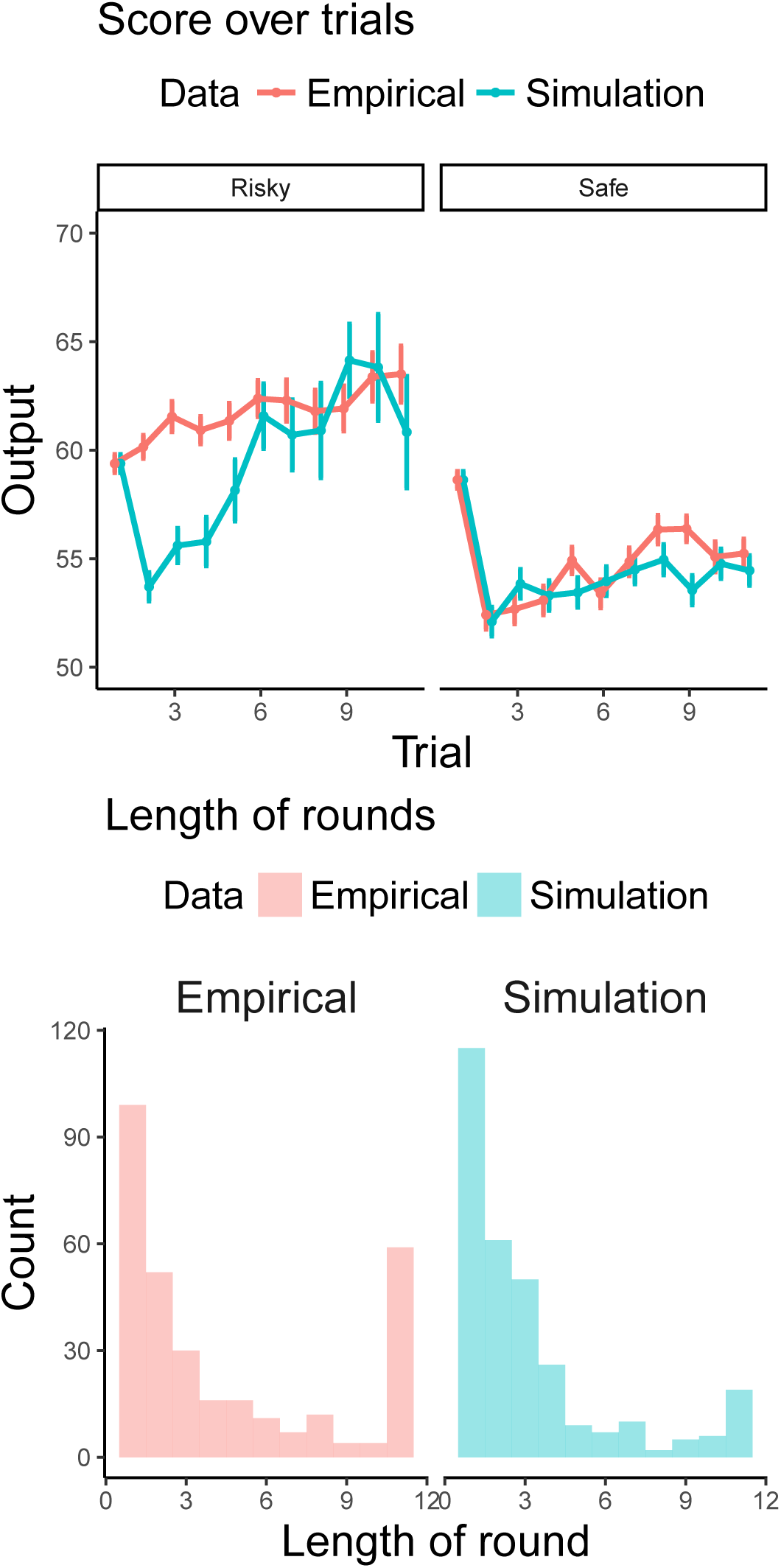
Posterior model checks of GP-POS model for Experiment 2. Upper panel shows the mean trajectories of scores over trials for both human participants and the GP-POS in the safe and risky rounds. Lower panel shows the histogram of lengths per round, i.e. how long both participants and the model managed to sample without sampling below the threshold.

### Discussion

Within a bivariate version of the risky spatially correlated multi-armed bandit, we found that participants improved over trials within the risky but not the safe condition, gained better scores than would be expected from random choices, played for longer than expected by chance in the risky conditions, and seemed to generalize further than in the previous task with univariate inputs. Our mixed effects regression analysis revealed that participants selected safe and maximizing points frequently, only marginally cared about choosing inputs from the expanding set input, and focused less on safe input points during safe conditions. The model comparison results showed that the Gaussian process regression model predicted behavior best in both conditions, even though predictions turned out to be generally better for the risky rounds than for the safe rounds. Importantly, whereas the different decision strategies predicted behavior equally well for the safe conditions, the probability of staying safe predicted behavior best for the risky condition. These results suggest that participants adapted their decision strategy to the task requirements while learning and generalizing about the underlying function in a similar fashion. In a posterior predictive check for the GP-POS model, we found that participants explored even more risk-aversely in the risky condition than predicted by the GP-POS model.

## General discussion

Learning unknown functions and exploiting this knowledge to maximize rewards are essential cognitive skills. Here we focused on a risky version of the spatially correlated bandit task, in which outcomes below a given threshold need to be avoided. We first analyzed participants’ choices using a rational Gaussian Process safe optimization strategy that establishes a safe set and tries to maximize outputs or expand the safe set by choosing inputs from this set. We found that participants shunned risks by focusing on maximizing outputs locally to “tried-and-tested” inputs, mostly ignoring lesser known input points which could potentially expand the safe set. This focus on avoiding unsafe inputs is consistent with a biological homeostasis maintenance principle (Korn & Bach, 2015, 2018) that prioritizes not losing everything over gaining maximum rewards (Houston, McNamara, & Hutchinson, 1993).

The results of our cross-validation model comparison revealed that participants learn and perform generalization in a similar fashion, in scenarios with and without risky constraints. While participants seemed to learn a similar representation of the reward function (using the same learning strategy) across the different task demands in Experiment 2, they did adapt their decision strategy to the riskiness of the environment, sometimes even more than predicted by the best currently available model. This in turn suggests a flexible mechanism that can guide people through risky environments via generalization and adaptive search.

In future work, we aim to focus on the factors which drive participants to switch from pure exploration to safe search strategies, and the situations in which switching constitutes a normative strategy, for example because it minimizes costs (Bach, 2015). Another promising avenue for future research will be marrying the powerful methods of generalization put forward here with restricted methods of planning that have been studied in other reinforcement learning tasks (Huys et al., 2015; Solway & Botvinick, 2015). Furthermore, investigating how different clinical populations differ in their search and generalization behavior when confronted with risky decision making tasks promises to extend our notion of computational mechanisms involved in mental illnesses (Huys, Maia, & Frank, 2016; Montague, Dolan, Friston, & Dayan, 2012).

Building and assessing additional sampling strategies with increased (and parametrically varying) levels of risk aversion as well as further probing the effect of different levels of riskiness on those strategies will also be two necessary steps for follow-up experiments.

### Related work

Unlike previous work on human behavior in a bandit setting, which has primarily focused on pure exploration and exploitation, our work addresses a relatively novel facet—optimizing risky functions while staying above a threshold. However, we note that this type of constrained risky choice situation, in which choices above a certain threshold or of a particular option abort the reward-accumulation process, has been investigated using other paradigms before.

One such task is the Balloon Analogue Risk Task (BART) in which participants can incrementally pump up a (digitally presented) balloon by clicking a button (Lejuez et al., 2002). On every trial, participants can gain more money by pumping the balloon to a larger size or gain nothing if the balloon explodes. A typical finding in studies using the BART is that people do not explore enough and behave relatively cautiously, a finding that aligns well with the results reported here. The Columbia Card Task is another similar paradigm, in which participants can turn around as many of 32 cards sequentially as they like as long as they encounter gains, however the trial is terminated as soon as a loss is encountered (Figner, Mackinlay, Wilkening, & Weber, 2009).

Another class of related tasks studies (optimal) foraging behavior in humans and other animals (Hills et al., 2015). Some of these studies share features with our proposed spatially correlated multi-armed bandit task such as a “clumpiness” of resources (Wilke et al., 2015) and a particular focus on model-based exploration (Kolling & Akam, 2017).

There have also been some more theoretical studies that assessed how participants solve the exploration-exploitation dilemma when rewards are correlated. For example, Reverdy, Srivastava, and Leonard (2014) and Reverdy and Leonard (2016) studied participants’ performance in a spatial multi-armed bandit problem, where Reverdy et al. (2014) developed a model of upper credible set sampling, whileReverdy and Leonard (2016) proposed a model fitting procedure and applied it to study differences in strategies adopted by subjects faced with different underlying environments.

### Conclusion

We introduced a novel paradigm to assess how participants search for spatially correlated rewards in scenarios where they have to avoid sampling below a threshold. Our results show that participants can adapt their sampling behavior to the underlying riskiness of the task, but explore only very cautiously overall. We expect that our approach of assessing safe optimization in humans will continue to enrich our understanding of how people resourcefully obtain rewards in the real world.

## Acknowledgments

We thank Dominik Bach and Felix Berkenkamp for helpful comments and discussions. E.S. was supported by a postdoctoral fellowship from the Harvard Data Science Initiative.

## Footnotes

1 Marr (1982) famously proposed to analyze intelligent systems on three different levels: the computational (what is the task the system is trying to solve), the algorithmic level (how does it solve it), and the implementation level (how is the solution implemented).

2 Sometimes the RBF kernel is specified as 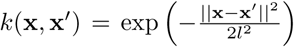 whereas we use *λ* = 2*l*^2^ for simplicity.

3 Although there are other ways according to which one could choose a particular value for *ω*, we follow standard practice in setting *ω* = 3 to ensure *high safety*.

